## Supplemental Material for "Time-of-day defines the efficacy of NAD^+^ to treat diet-induced metabolic disease by adjusting oscillations of the hepatic circadian clock"

### Supplementary Figure legends

**Figure S1. Chronic 50 mg/Kg IP injection of $NAD^+$ at ZT11 recovers daily rhythms in hepatic $NAD^+$ in obese mice**

(**A**) $NAD^+$ was IP injected in mice at the indicated doses. One hour later, mice were sacrificed, and livers were snap frozen in liquid nitrogen. Hepatic $NAD^+$ was quantified by HLPC (n = 3 mice and 3 technical replicates per dose). (**B**) Energy intake and relative energy intake per week (n = 20 mice per group) (**C**) Energy intake throughout the day and relative energy intake per day (n = 10 mice per group). (**D**) Total energy intake during day and night periods (n = 10 mice per group). (**E**) Daily rhythms in hepatic $NAD^+$ was evaluated by CircWave. (n = 5 mice per time point and 3 technical replicates) (**F**) Rhythmic insulin levels evaluated with CircWave (**G**) Circadian rectal temperature before $NAD^+$ treatment (n = 10 mice per group and 3 technical replicates) (**H**) Representative thermography image at ZT12 at week 8, before $NAD^+$ treatment (n = 4 mice per group). (**I**) Circadian rectal temperature 20 days after starting with $NAD^+$ treatment at ZT11 (n = 10 mice per group and 3 technical replicates) (**J**) Representative thermography image at ZT12, after 20 days on $NAD^+$ treatment at ZT11 (n = 4).(**K-N**) Glucose (K,L) and insulin (M,N) tolerance tests were performed at both the rest (K, M; ZT4) and the active (L, N; ZT16) period the day before $NAD^+$ treatments (n = 5-6 mice). Here, mice within HFN group had not been treated yet. AUC: area under the curve.

CD: Control diet fed mice; HF: High-fat diet fed mice; HFN: High-fat diet fed, $NAD^+$ treated mice at ZT11. Data represent mean ± SE and were analyzed by two-way ANOVA using Tukey posttest, except for bar graphs, where one-way ANOVA followed by Tukey's posttest was used. Circadian rhythmicity was assessed by CircWave. * $p$ <0.05, ** $p$ <0.01, *** $p$ <0.001. Symbol key for comparisons: * CD vs HF; + CD vs HFN; # HF vs HFN. Data from live mice were replicated in two independent experiments.

**Figure S2. Circadian rhythms in hepatic triglycerides**

(**A**) CircWave results for the investigations of circadian rhythmicity in hepatic triglycerides. CD: Control diet fed mice; HF: High-fat diet fed mice; HFN: High-fat diet fed, $NAD^+$ treated mice at ZT11.. Circadian rhythmicity was confirmed when the CircWave *F* test produced a significant value ($P<0.05$).

**Figure S3. Immune response and energy sensing pathways are corrected by a $NAD^+$ chronotherapy in obese mice.**

(**A, C**) Gene set enrichment analysis (GSEA) investigated within the molecular signature database (MSigDB) "Hallmark" gene set collection. Genes were rank-ordered by differential expression (DE) between obese mice untreated (HF) or treated with timed $NAD^+$ therapy at ZT11 (HFN), specifically at the rest phase (A; ZT6) or at the active phase (C; ZT18) (**B, D**) Heatmap depicting genes pertaining to the indicated gene set from the MSigDB, and rank-ordered according to their DE between HF and HFN groups, at ZT6 (B) or at ZT18 (D). (**E**) Heatmap illustrating changes in expression from known target genes of FOXA2 (HNF3-β) transcription factor. $NAD^+$ chronotherapy elicits significant changes in expression from these genes. NES, normalized enrichment score; FDR, false discovery rate-adjusted *q* value.

**Figure S4. Metabolic sensors regulated by $NAD^+$ chronotherapy**

(**A**, **B**) Quantification of western blots from n = 4 – 5 mice for the indicated proteins and relative phosphoprotein levels. All measurements were normalized to the tubulin loading control, and data from CD at ZT0 was set to 1 (**C**) Western blot for ULK1 and p-ULK1(S555) proteins from liver whole cell extracts sampled at ZT12 (left), and corresponding quantification by densitometry (right). Tubulin was used as loading control (**D**-**H**) Quantification of western blots from n = 3 – 5 mice for the indicated proteins and relative phosphoprotein levels. All measurements were normalized to their corresponding tubulin or p84 loading controls, and data from CD at ZT0 was set to 1.

CD: Control diet fed mice; HF: High-fat diet fed mice; HFN: High-fat diet fed, $NAD^+$ treated mice at ZT11. Points at ZT24 are duplicates of ZT0 replotted to show 24-h trends. Data represent mean ± SE and were analyzed by two-way ANOVA using Tukey posttest, except for bar graphs, where one-way ANOVA followed by Tukey's posttest was used. * p <0.05, ** p <0.01, *** p <0.001. Symbol key for comparisons: $^{*}$ CD vs HF; $^{+}$ CD vs HFN; $^{\#}$ HF vs HFN.

**Figure S5. Neutral lipid accumulation in the liver is relieved by NAD$^+$ in obese mice independently of time of treatment.**

(**A**) Hepatic NAD$^+$ content measured by HPLC along the day at the indicated times for all groups after the experimental paradigm (n = 4 biological replicates, 3 technical replicates) (**B**) Circadian rhythms in hepatic NAD$^+$ from mice treated with NAD+ at ZT23 (HFN23) was evaluated by CircWave. (n = 4-5 mice per time point and 3 technical replicates) (**C**) Three weeks of food intake before and after the treatment were averaged for n= HF: 48; HFN: 37 ; HFN23: 27. Two-way ANOVA with Bonferroni post-test was applied, and the data are means ± SD (**D**) Circadian serum levels of insulin (n = 5 mice per time point and 2 technical replicates). AUC: area under the curve. (**E**) Circulating glucose levels were measured in fasted mice at ZT4 and at ZT16 (n=9-17). (**F**, **G**) Glucose (F) and insulin (G) tolerance tests were performed at ZT4, on the day before NAD$^+$ treatments (day0), after 10 days of treatment (day 10) and at the end of the treatment (day20) (n = 5-6 mice). AUC: area under the curve. (**H**) Representative hepatic histopathology. Upper panel: Oil-red-O stain (ORO). Lower panel: Hematoxilin/Eosin. Images were acquired at 20X optical magnification, and detailed 100X magnification is shown (**I**) Bar graph represents quantification of ORO signal in arbitrary units. Signal for control mice was set to 1 (n = 3 biological and 3 technical replicates). (**I**) The length of lipid droplets was compared between groups (n = 3 biological and 3 technical replicates).

CD: Control diet fed mice; HF: High-fat diet fed mice; HFN: High-fat diet fed, NAD$^+$ treated mice at ZT11; HFN23: High-fat diet fed, NAD$^+$ treated mice at ZT23. Points at ZT24 are duplicates of ZT0 replotted to show 24-h trends Data represent mean ± SE and were analyzed by one-way ANOVA followed by Tukey's posttest. * p <0.05, ** p <0.01, *** p <0.001.

**Figure S6. Daily levels of nutrient sensors and gene expression profiles in the liver.**

(**A**) Quantification of western blots from liver (n = 4 – 5 mice) for the indicated proteins across the day. All measurements were normalized to their corresponding GAPDH or p84 loading control, and data from CD at ZT0 was set to 1. (**B**) Quantification by densitometry from western blots for RSK protein in liver whole cell extracts (n = 4). Student´s *t* test; n.s: non-significant (**C**) RT-qPCR expression data in the liver for the indicated genes at ZT18 (n= 5-6 biological replicates per data point) (**D**) RT-qPCR was used to assess hepatic expression of housekeeping genes across the day (n = 4-6 biological replicates per data point) (**E**) RT-qPCR was used to determine expression of anti-obesity genes in the liver form mice at the indicated times-of-day (ZT) (n = 5-6 biological replicates per data point).

The data are means ± SE. *p < 0.05, **p < 0.01, ***p < 0.001, Two-way ANOVA followed by Tukey's post test. Points at ZT24 are duplicates of ZT0 replotted to show 24-h trends. Symbol key for multiple comparisons: $^{*}$CD vs HF, $^{+}$ CD vs HFN, $^{\circ}$CD vs HFN23, $^{\#}$ HF vs HFN, ; $^{x}$ HF vs HFN23, $ HFN vs HFN23.

**Figure S7. Time-of-day dependent effects of NAD$^{+}$ supply on circadian gene expression.**

(**A**) Acrophase and *p*-value for circadian rhythmicity from clock and clock-controlled gene expression was assessed by CircaWave. (**B**) RT-qPCR was used to determine expression of genes related to lipid metabolism in the liver form mice at the indicated times-of-day (ZT) (n = 5-6 biological replicates per data point). (**C**) Total cholesterol was measured in livers from the indicated groups of mice, at selected ZTs (n= 4-5 mice per data point). Mg of cholesterol per gram of liver are plotted. (**D**) CircaWave analyses of rhythmicity from the hepatic cholesterol data. (**E**) Average 24-hour activity profile from the indicated groups of mice. Average was calculated for five days before NAD$^{+}$ treatment. n= 6-7 mice. CD: Control diet fed mice; HF: High-fat diet fed mice; HFN: High-fat diet fed, NAD+ treated mice at ZT11; HFN23: High-fat diet fed, NAD+ treated mice at ZT23. Points at ZT24 are duplicates of ZT0 replotted to show 24-h trends Data represent mean ± SE

**Figure S8. Time of day-dependent effects of the NAD$^{+}$ precursor nicotinamide (NAM) to treat diet-induced obesity.**

(**A**) Mice fed a high-fat diet were IP injected at week 8, with 200 mg/Kg of NAM either at ZT11 (HF_NAM11) or at ZT23 (HF_NAM23) for three weeks. Weekly body weight measurements are shown (n = 4 mice per group). Red arrow indicates the period of treatment with NAM. (**B**) Percent change in body weight between weeks 8 (just before treatment), and 11 (end of the treatment). Two tailed t-test (**C**, **D**) Glucose (GTT) and insulin (ITT) tolerance tests were performed at ZT4 before (day 0) and after (day 20) NAM treatments (n = 4 mice). AUC: Area under de curve. Two tailed t-test. (**E**) Clock protein expression at ZT12 was measured by western blot from liver whole cell extracts of obese mice treated with NAM at ZT11 (HF_NAM11) or ZT23 (HF_NAM23). Tubulin or GAPDH were used as loading control. Quantification was performed from n=3 mice. Two tailed t-test (**F**) Representative double plotted actograms of locomotion measured using infrared sensors in a 12- hour light/12-hour dark cycle. The data represent means ± SE. *p < 0.05, **p < 0.01, ***p < 0.001.

**Supplementary Tables**

**Table S1**: List of DE genes between ZT6 and Z18, and their biological processes

**Table S2**: List of common genes in comparisons SD-HF vs HF-HFN and their functional analyses.

**Table S3**: List of common genes in comparisons SD-HFN vs HF-HFN and their functional analyses

**Table S4**: List of primers used in this study

Escalante-Covarrubias *et al.* **Supplementary Figure S1**

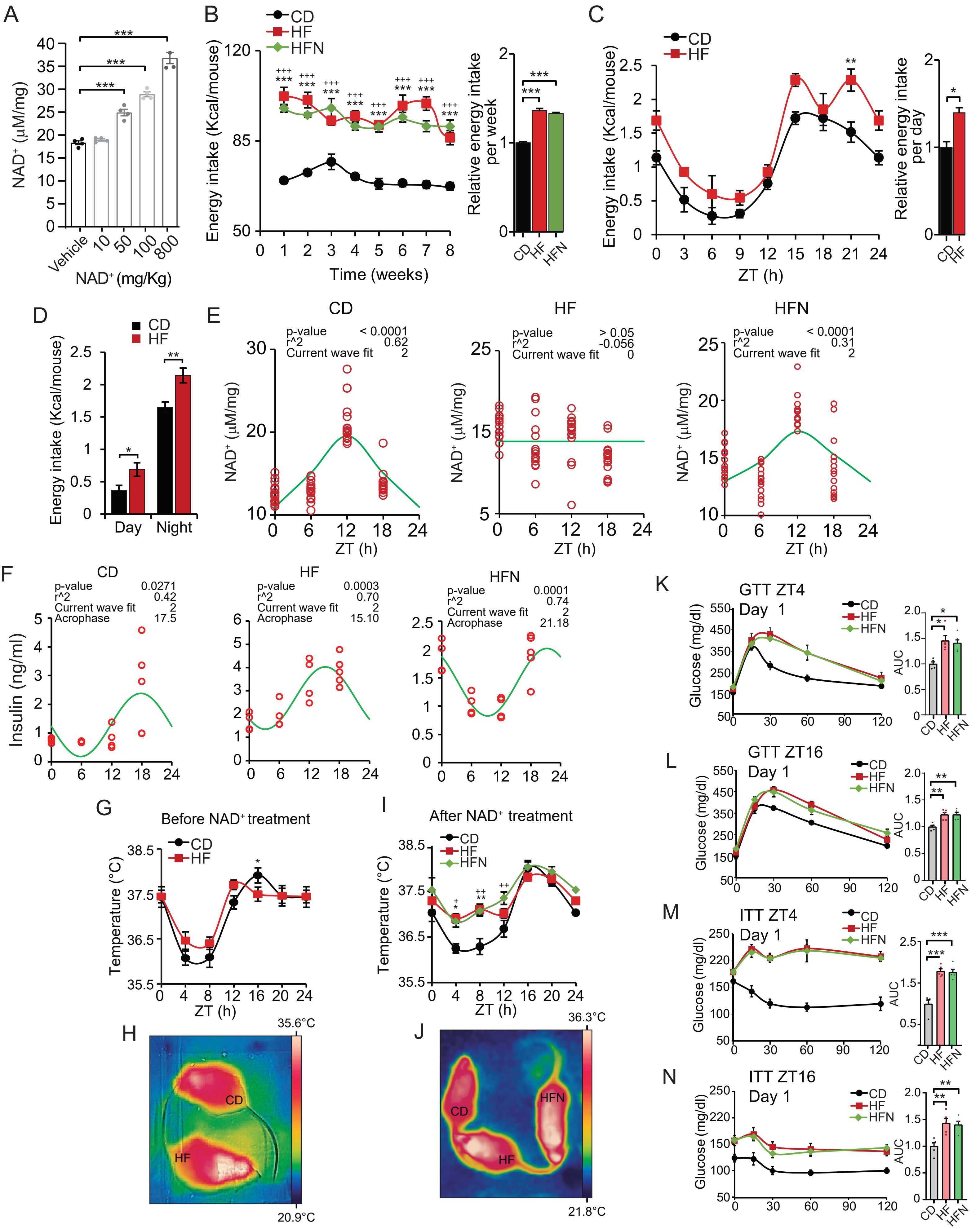

Escalante-Covarrubias *et al.* **Supplementary Figure S2**

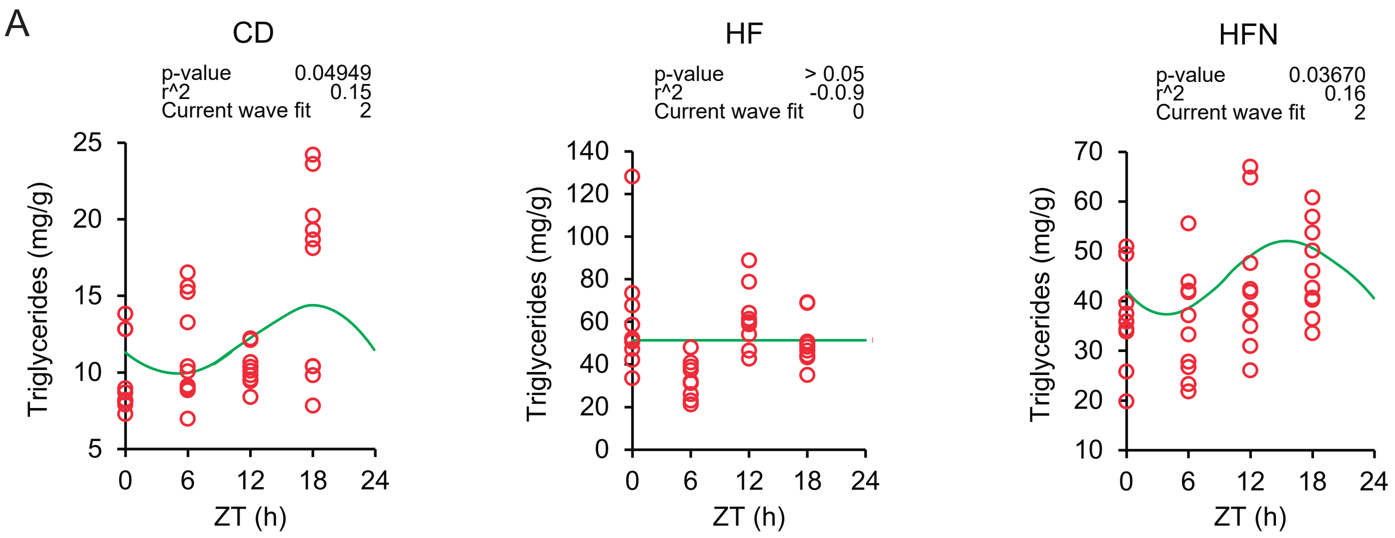

Escalante-Covarrubias *et al.* **Supplementary Figure S3**

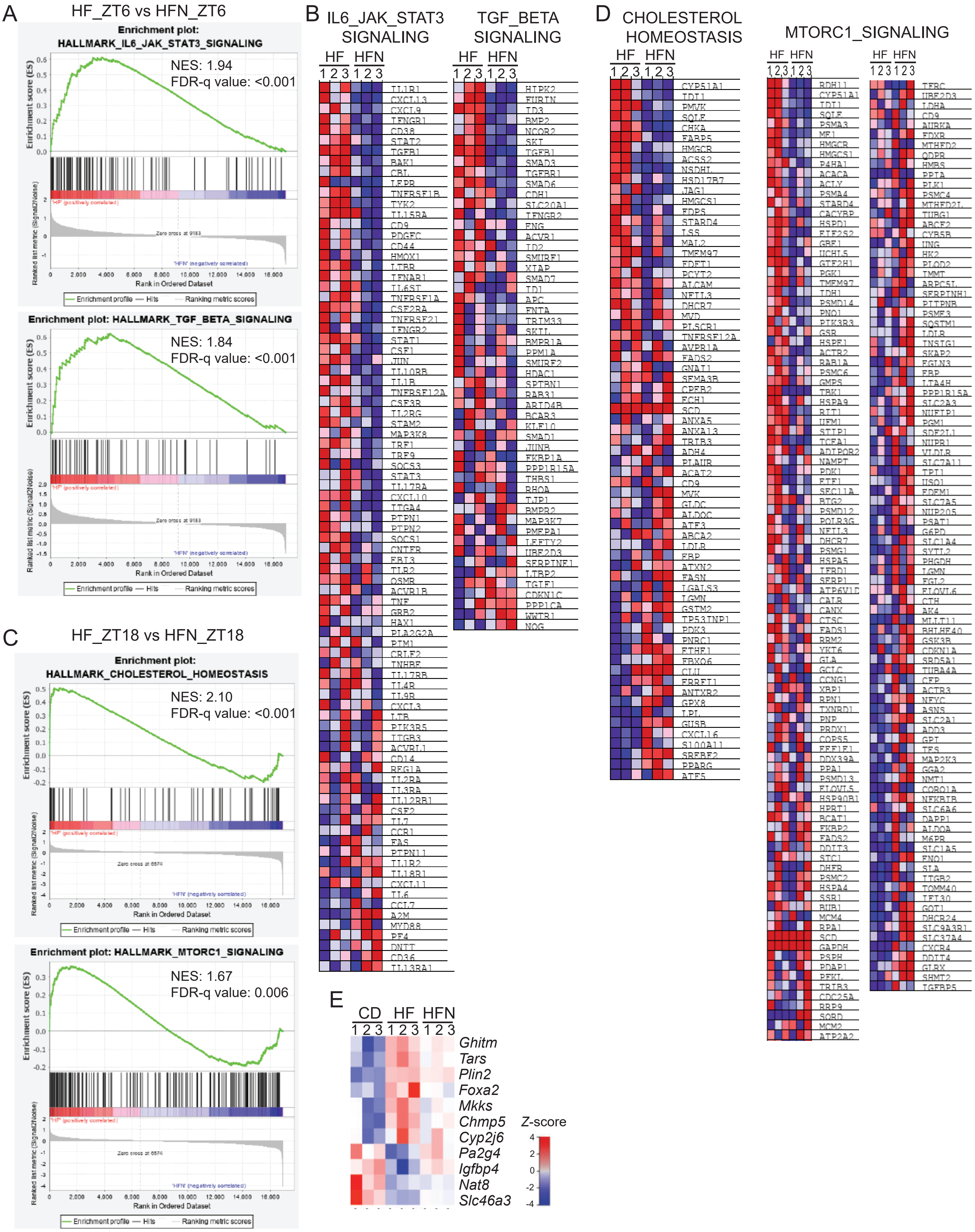

Escalante-Covarrubias *et al.* **Supplementary Figure S4**

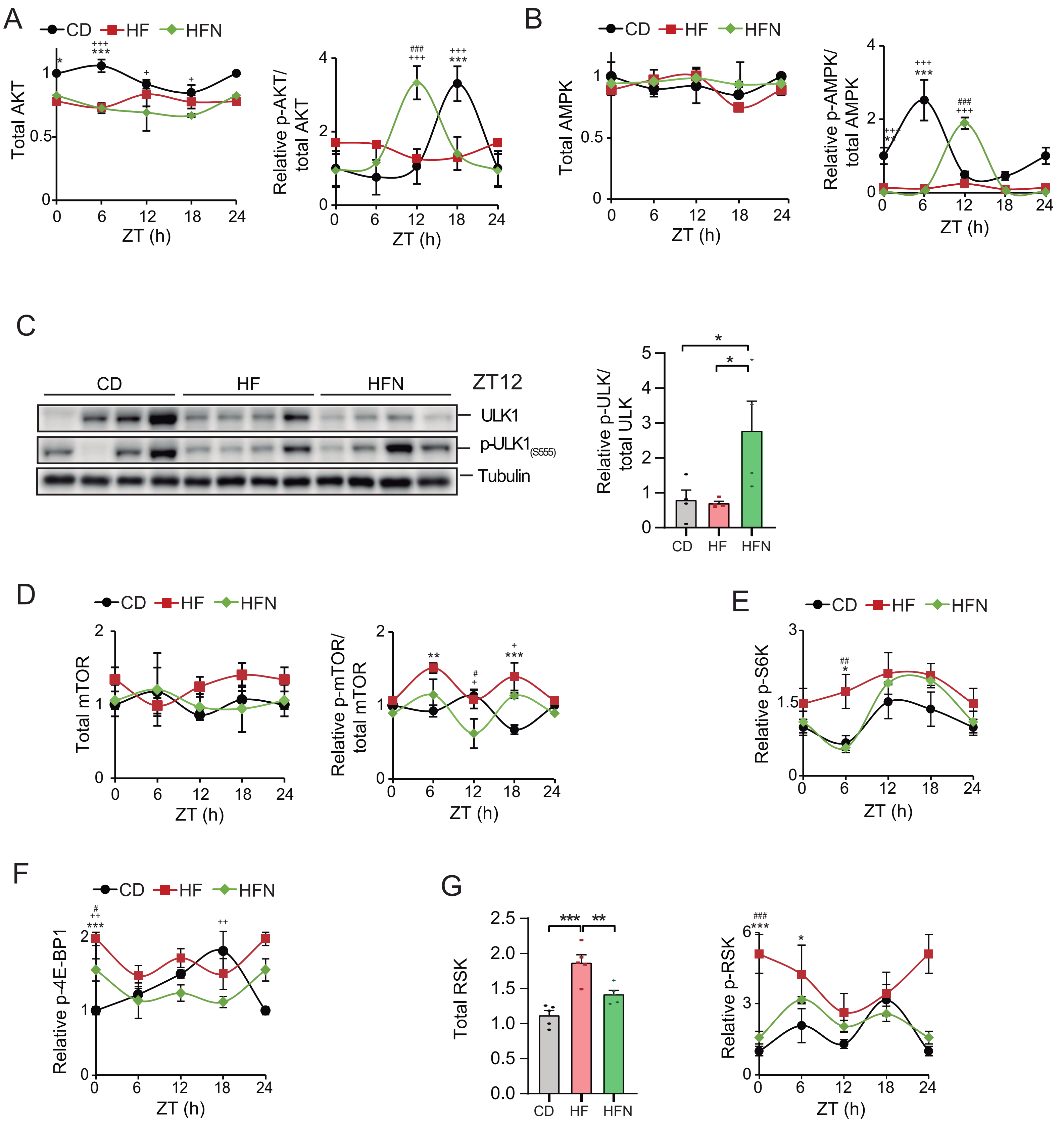

Escalante-Covarrubias *et al.* **Supplementary Figure S5**

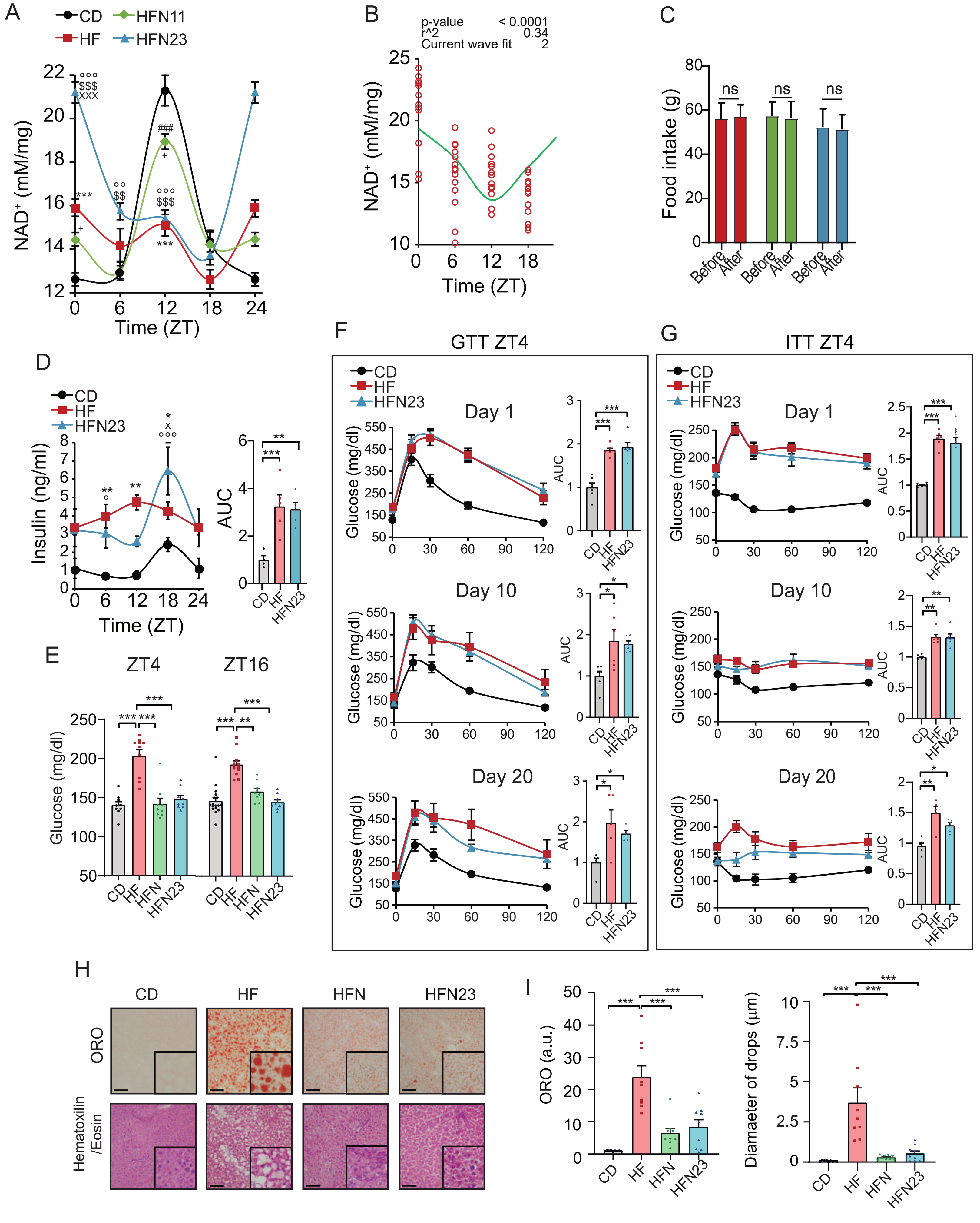

Escalante-Covarrubias *et al.* **Supplementary Figure S6**

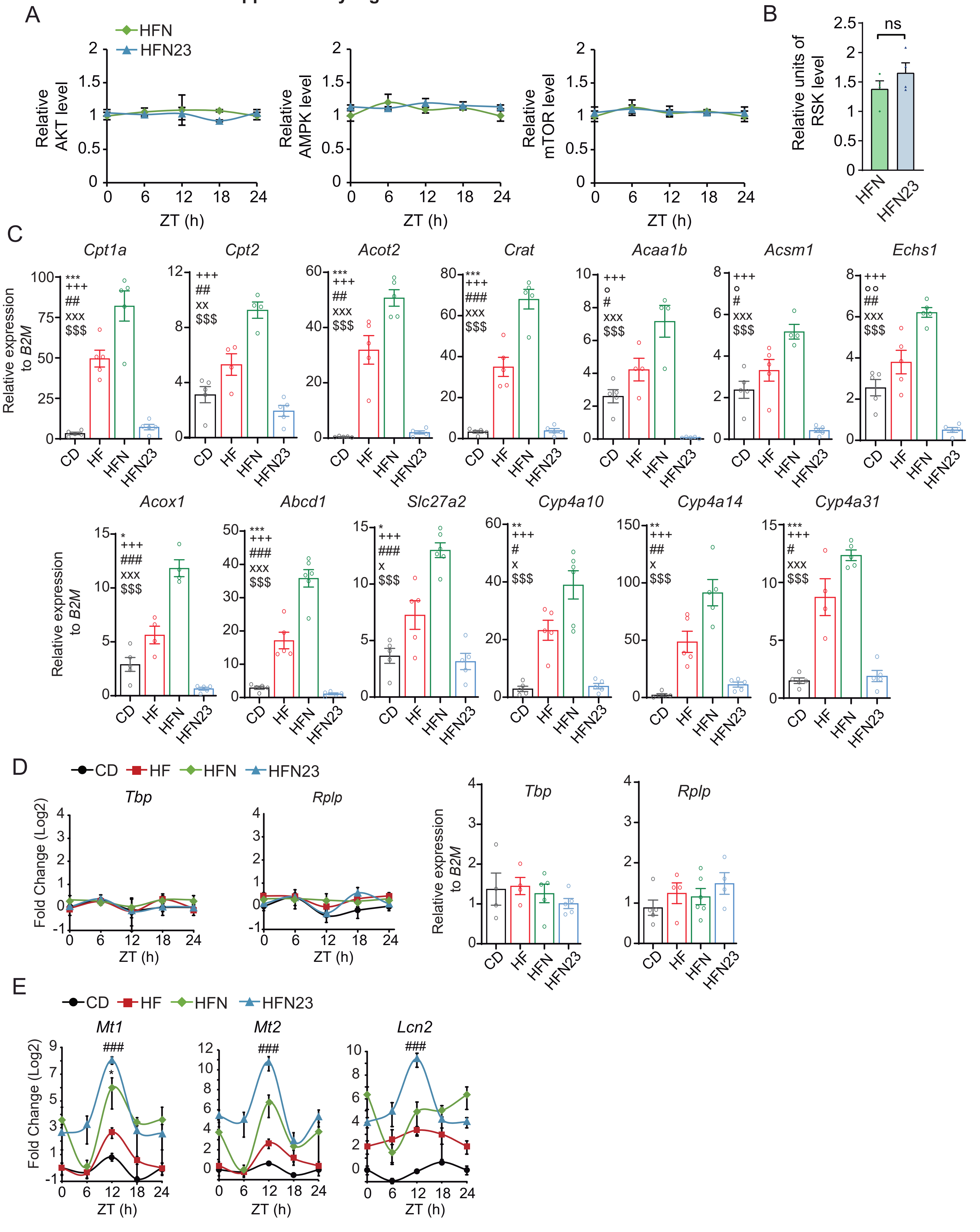

A

| | | Acrophase (hour) | *P*-value |
|---|---|---|---|
| *Bmal1* | CD | 18.771 | 0.000112 |
| | HF | 18.659 | 0.000031 |
| | HFN | 18.507 | 0.000047 |
| | HFN23 | 8.6 | 0.000005 |
| *Clock* | CD | 18.25 | 0.0094 |
| | HF | 18.34 | 0.000039 |
| | HFN | 17.74 | 0.0012 |
| | HFN23 | 5.97 | 0.013 |
| *RevErbα* | CD | 6.89 | 0.000008 |
| | HF | 5.39 | 0.000001 |
| | HFN | 6.07 | 0.000021 |
| | HFN23 | 18.98 | 0.0047 |
| *Cry1* | CD | 17.59 | 0.004 |
| | HF | 16.85 | 0.000108 |
| | HFN | 17.29 | 0.000062 |
| | HFN23 | 6.57 | 0.0026 |
| *Per1* | CD | 13.26 | 0.03 |
| | HF | 11.35 | 0.0048 |
| | HFN | 12.95 | 0.000002 |
| | HFN23 | 1.707 | 0.00032 |
| *Per2* | CD | 14.79 | 0.0025 |
| | HF | 13.9 | 0.000009 |
| | HFN | 14.63 | 0.00016 |
| | HFN23 | 3.43 | 0.005 |

| | | Acrophase (hour) | *P*-value |
|---|---|---|---|
| *Dbp* | CD | 8.77 | 0.000051 |
| | HF | 7.08 | 0.000029 |
| | HFN | 7.56 | 0.000008 |
| | HFN23 | 22.77 | 0 |
| *Tef* | CD | 14.34 | 0.018 |
| | HF | 11.94 | 0.0025 |
| | HFN | 13.46 | 0.0021 |
| | HFN23 | 0.93 | 0.000135 |
| *Nfil3* | CD | 18.36 | 0.002 |
| | HF | 18.85 | 0.000001 |
| | HFN | 17.8 | 0.00015 |
| | HFN23 | 10.4 | 0.026746 |
| *Noct* | CD | 14.1 | 0.028 |
| | HF | 11.82 | 0.014 |
| | HFN | 12.74 | 0.000001 |
| | HFN23 | 0.08 | 0.000201 |

B

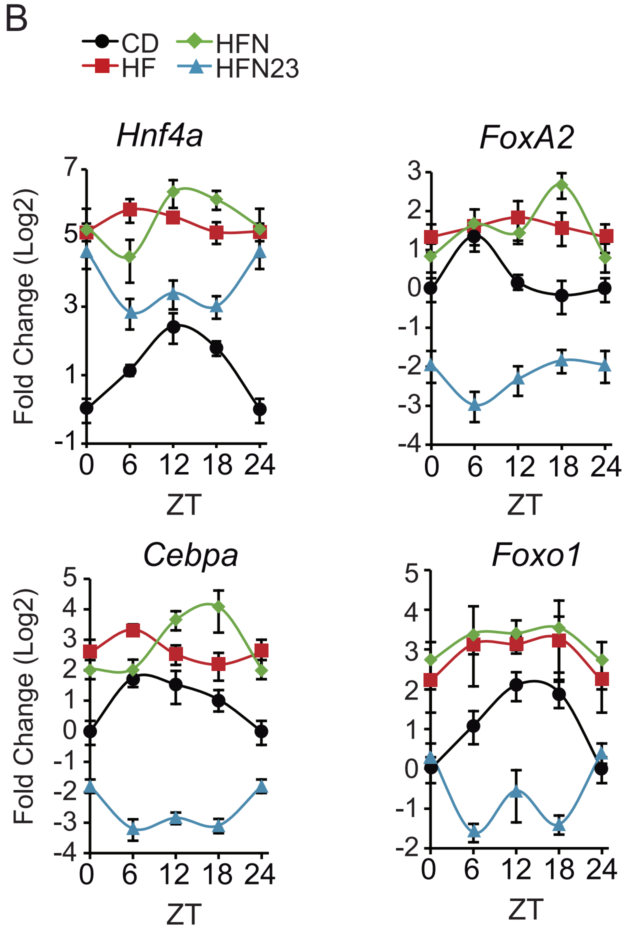


C

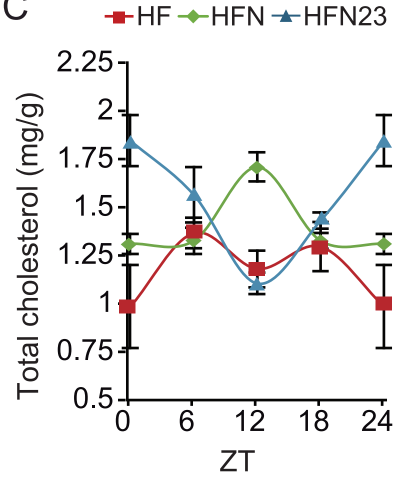


D

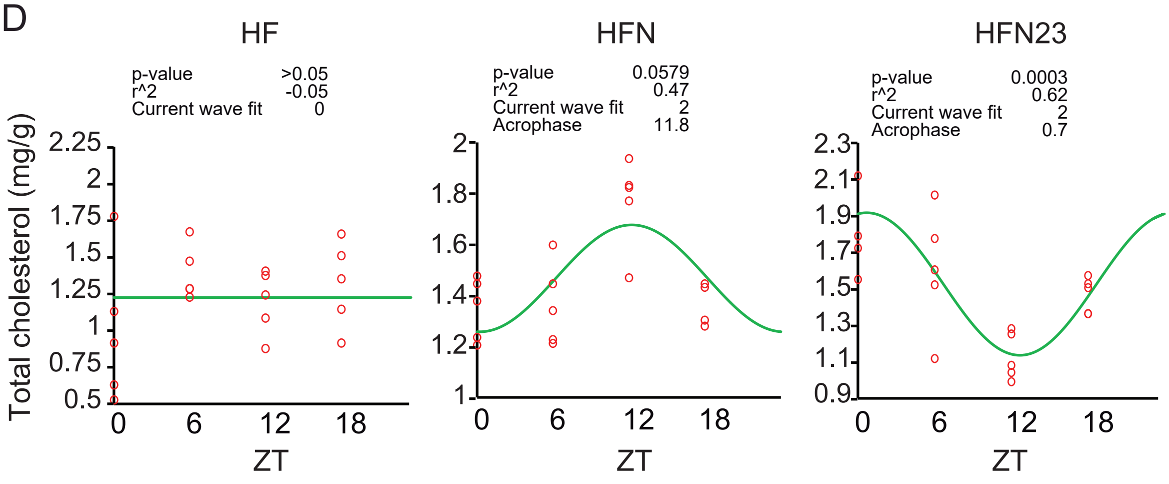


E

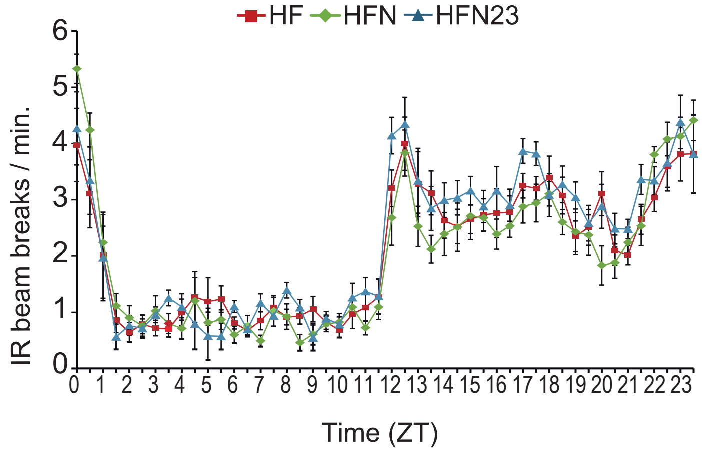

Escalante-Covarrubias *et al.* **Supplementary Figure S8**

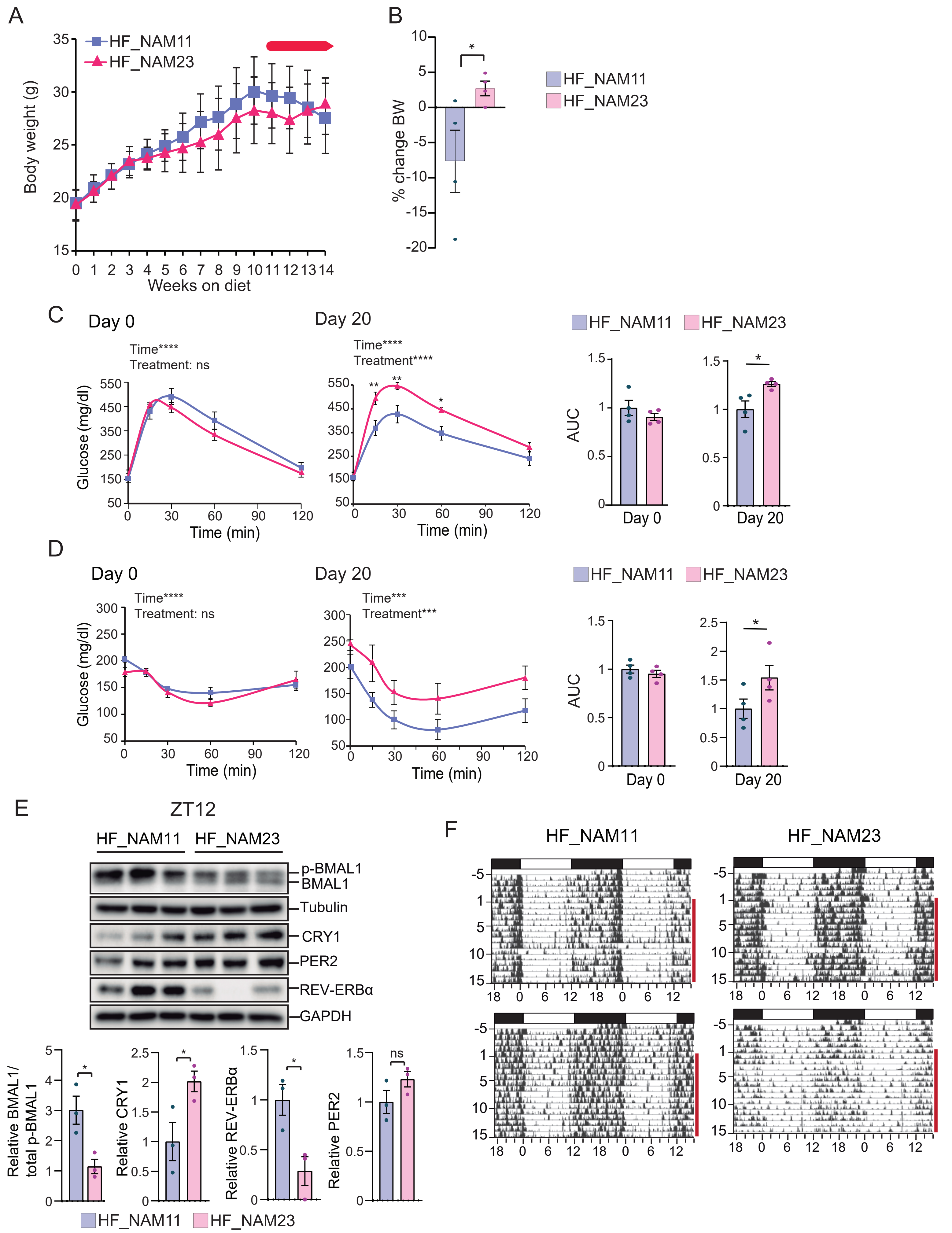

**mRNA primers**

| Primer | Sequence |
|---|---|
| ***Abcd1*** *Fw* | GCC AGG GTG TAC GAG ATG |
| ***Abcd1*** *Rv* | CCT CTA CAT GGA CAC CAG ACT G |
| ***Acaa1b*** *Fw* | CA TGC TGA GAT TGT GCC TGT G |
| ***Acaa1b*** *Rv* | GGT AGA GCC TCC ATC CTT GAA G |
| ***Acot2*** *Fw* | CTG AGA GCA AGC AGG TTG TG |
| ***Acot2*** *Rv* | GCT CAG CGT CGC ATT TGT C |
| ***Acox1*** *Fw* | CAT CAC AGG GGC TCA GAT GTC |
| ***Acox1*** *Rv* | CAG TGG GGA CTT CTT GGC |
| ***Acsm1*** *Fw* | CCT TGA TTC TGC CCC GAG TG |
| ***Acsm1*** *Rv* | GTC CTT GGC TTT CAG TTG GGT |
| ***B2m*** *Fw* | GGT CTT TCT GGT GCT TGT CTC A |
| ***B2m*** *Rv* | GTT CGG CTT CCC ATT CTC C |
| ***Bmal1*** *Fw* | CCA AGA AAG TAT GGA CAC AGA CAA A |
| ***Bmal1*** *Rv* | GCA TTC TTG ATC CTT CCT TGG T |
| **Clock** Fw | ACCACAGCAACAGCAACAAC |
| **Clock** Rv | GGCTGCTGAACTGAAGGAAG |
| ***Cpt1a*** *Fw* | CCA TGA AGC CCT CAA ACA GAT C |
| ***Cpt1a*** *Rv* | ATC ACA CCC ACC ACC ACG ATA |
| ***Cpt2*** *Fw* | CAG AAG CCT CTC TTG AAT GAC AGC |
| ***Cpt2*** *Rv* | GCA GCT CCT TCC CAA TGC |
| ***Crat*** *Fw* | AAG AAT GGG CTC ACA CCA AG |
| ***Crat*** *Rv* | CGC TCC AGT CCC TTC TGT AG |
| **Cry1** Fw | CTGGCGTGGAAGTCATCGT |
| **Cry1** Rv | CTGTCCGCCATTGAGTTCTATG |
| ***Cyp4a10*** *Fw* | ACT TCC CAA GTG CCT TTC C |
| ***Cyp4a10*** *Rv* | TAC GCA CCA TTA GCC TTT GG |
| ***Cyp4a14*** *Fw* | CAA GGC AGT GTT CAG TTG GAT G |
| ***Cyp4a14*** *Rv* | CAG GCG AAA GAA AGT CAG GTT G |
| ***Cyp4a31*** *Fw* | CCT CTG TGT TCT GTC TGC TCC |
| ***Cyp4a31*** *Rv* | GTG AAA GGG CGG TGA TGG |
| ***Dbp*** *Fw* | GCT CCA GTA CTT CTC ATC CTT CTG T |
| ***Dbp*** *Rv* | AAT GAC CTT TGA ACC TGA TCC CGC T |
| ***Echs1*** *Fw* | GGG CTG ATC CAG TTG AAC C |
| ***Echs1*** *Rv* | CAC CCA CAG CAG GAT CTT G |
| ***Lcn2*** *Fw* | CAG AAG GCA GCT TTA CGA TGT AC |
| ***Lcn2*** *Rv* | TCT GAT CCA GTA GCG ACA GC |
| ***Mt1*** *Fw* | ATG GA CCC CAA CTG CTC CTG |
| ***Mt1*** *Rv* | CCC TGG GCA CAT TTG GAG C |
| ***Mt2*** *Fw* | GCA AAG AGG CTT CCG ACA AG |
| ***Mt2*** *Rv* | AGG CTA GGC TTC TAC ATG GTC T |
| ***mtCOX1*** *Fw* | ACC ATC ATT TCT CCT TCT CCT A |
| ***mtCOX1*** *Rv* | TAG ATT TCC GGC TAG AGG TG |
| ***Nfil3*** *Fw* | GGTTACAGCCGCCCTTTCTT |

**ChIP primers**

| Primer | Sequence |
|---|---|
| Per2 Ebox for | TGCCACCTCATTTGCATACTG |
| Per2 Ebox rev | CGCAGCATCTTCATTGAGGA |
| Nr1d1 Ebox for | CACTCTGCCAATCTCAACCG |
| Nr1d1 Ebox rev | GCTCCCTGGAATCACATGGT |
| Cry1 Ebox for | CGGCACCTCACGTTTCTG |
| Cry1 Ebox rev | GGATCCCACGCGAGAACT |
| Tef I1 Ebox for | CCTCTGAGGTGGCTTCTCC |
| Tef I1 Ebox rev | CCCTTTCGCCACCACTTTC |
| Dbp I1 Ebox Fw | ATGCTCACACGGTGCAGACA |
| Dbp I1 Ebox Rv | CTGCTCAGGCACATTCCTCAT |
| Dbp 3´UTR Fw | GCCTGGAATGTATGAGCTAGCA |
| Dbp 3´UTR Rv | GGCACCGGAGTAGGCAAGA |
| Nmrk1 I1 Fw | TACAGGGTCAGCACAATGGT |
| Nmrk1 I1 Rv | TGCTCAGGTCATAGGTCTCC |
| Nampt Ebox Fw | GAGAAAAGCAAGTCCACGCG |
| Nampt Ebox Rv | TACCTTTGTCTCCCGCTTGG |
| Nmnat3 E1 Ebox for | TGCAGCACGTTTACAGTCAG |
| Nmnat3 E1 Ebox rev | TACACGTGACAATCGCTGG |
| Nadk I1 Ebox for | AGGACACAGCCGATATCAGG |
| Nadk I1 Ebox rev | TCGGTAACAAGTGTCCAGC |
| Ppara TSS for | CAGTGAGGTGGGTGGACAG |
| Ppara TSS rev | GGCCCTGAAAACTCGCAC |
| Pparg2 I1 Ebox for | CTCCCACGTTAGCAGTTTGG |
| Pparg2 I1 Ebox rev | GTTGGCAAGGAATTGTGGT |
| Srebp1c I1 Fw | CAGAGGGAAAGCAGAGGATGTGG |
| Srebp1c I1 Rv | CCACCTTGGGCTGGAAATTGGT |

| | |
|---|---|
| ***Nfil3*** *Rv* | TCCGGCACAGGGTAA ATCTG |
| ***Noct*** *Fw* | CACTCTCCCATTCGCGTCAT |
| ***Noct*** *Rv* | AGGCACTTCCTCTCTTCCCA |
| ***Per1*** *Fw* | GTG CAC AGC ACC CAG TTC CC |
| ***Per1*** *Rv* | ACC AGC GTG TCA TGA TGA CAT A |
| ***Per2*** Fw | GGCTTCACCATGCCTGTTGT |
| ***Per2*** Rv | GGAGTTATTTCGGAGGCAAGTGT |
| ***Rev-Erbα*** *Fw* | GGG CAC AAG CAA CAT TAC CA |
| ***Rev-Erbα*** *Rv* | CAC GTC CCC ACA CAC CTT AC |
| ***Rplp1*** *Fw* | GTG GTG CTG CTC CAT CC |
| ***Rplp1*** *Rv* | GGT TCA GCT CTT TAT TAG CCA ACT TA |
| ***S18*** *Fw* | TGG CTC ATT AAA TCA GTT ATG GT |
| ***S18*** *Rv* | GTC GGC ATG TAT TAG CTC TAG |
| ***Slc27a2*** *Fw* | ATG CCG TGT CCG TCT TTT AC |
| ***Slc27a2*** *Rv* | CTT CAG ACC TCC ACG ACT CC |
| ***Tbp*** *Fw* | CAA ACC CAG AAT TGT TCT CCT T |
| ***Tbp*** *Rv* | ATG TGG TCT TCC TGA ATC CCT |
| ***Tef*** *Fw* | TCTTTCAGCCCTCGGAAACC |
| ***Tef*** *Rv* | GCAGGGTCAGGGTTGAAGTT |
| ***Nmrk1*** *Fw* | TCGAACCCGCTTCGGATA |
| ***Nmrk1*** *Rv* | TGTCGTCTTCCCTCCGTTTG |
| ***Nampt*** *Fw* | GGTCATCTCCCGATTGAAGT |
| ***Nampt*** *Rv* | TCAATCCAATTGGTAAGCCA |
| ***Nmnat3*** *Fw* | CCGTCATCACCTACATCAGG |
| ***Nmnat3*** *Rv* | GCCAGTCTTTCCTTTCCCT |
| ***Nadk*** *Fw* | ATGCCTCCTCACTTTTCCAG |
| ***Nadk*** *Rv* | GAAAGCCCAGGGATCCAA |
| ***Pparg2*** *Fw* | TGGGTGAAACTCTGGGAGATTC |
| ***Pparg2*** *Rv* | AGAGGTCCACAGAGCTGATTCC |
| ***Ppara*** *Fw* | ACAAGGCCTCAGGGTACCA |
| ***Ppara*** *Rv* | GCCGAAAGAAGCCCTTACAG |
| ***Srebf1c*** *Fw* | GGAGCCATGGATTGCACATT |
| ***Srebf1c*** *Rv* | GGCCCGGGAAGTCACTGT |
| ***Hnf4a*** *FW* | GTTCTGTCCCAGCAGATCAC |
| ***Hnf4a*** *Rv* | GCTCCTTCATAGACTCACACAC |
| ***FoxA2*** *Fw* | GAGCACCATTACGCCTTCAAC |
| ***FoxA2*** *Rv* | TGCATGACCTGTTCGTAGGC |
| ***Cebpa*** *Fw* | GGTGCGTCTAAGATGAGGGA |
| ***Cebpa*** *Rv* | AATACTAGTACTGCCGGGCC |
| ***Foxo1*** *Fw* | ACGAGTGGATGGTGAAGAGC |
| ***Foxo1*** *Rv* | TGCTGTGAAGGGACAGATTG |
